# Cytochrome *c* as a facultative enzyme – lipoperoxidase and it’s inhibition by antioxidants

**DOI:** 10.1101/838540

**Authors:** L. A. Romodin, Y. A. Vladimirov, N. P. Lysenko, V. B. Chernetsov, Y. V. Antonova

## Abstract

Many researchers consider a key role in initiation of apoptosis along the mitochondrial pathway to be enhanced by cytochrome c, one of the components of the mitochondrial respiratory chain, which acquires peroxidase activity by forming a complex with phospholipids. Mitochondrial membranes are destroyed affected by the peroxidase reaction catalyzed by this supramolecular nanoparticle, resulting in the release of various proapoptotic factors into the cellular cytoplasm, ultimately leading to the development of an apoptosis pathway. The study of lipoperoxidase activity of the cytochrome c with cardiolipin complex is conducted via activated chemiluminescence. However, prior to this study, no assessment of the potential contribution of free non-heme iron, which can be inserted into the sample, into chemiluminescence of the system of cytochrome c complex with cardiolipin– hydrogen peroxide. It was found during the study process, that chemiluminescence of this system is indeed generated by the activity of the cytochrome c with cardiolipin complex, and the method of activated chemiluminescence is actually suitable for its study. The effect of trolox and dihydroquercetin (taxifolin) as synthetic and natural antioxidants on lipoperoxidase activity of the cytochrome c with cardiolipin complex was as well assessed via application of chemiluminescence activator specific for lipid peroxidation reactions – coumarin-334. A complete inhibition of lipoperoxidase activity for a few minutes with its subsequent full development under the trolox response and its dose-dependent uniform decrease under dihydroquercetin effect was obtained. These findings are promising for the future studies on inhibition of lipoperoxidase activity of this nanoparticle by antioxidants in order to inhibit the inappropriate apoptosis. Peroxidase activity of intact mitochondria in the comparative application of two chemiluminescence activators: coumarin-334 and coumarin-525, was also featured.

**Significance of the study:** This study proves that the method of activated chemiluminescence is adequate to study the processes occurring in the early stages of apoptosis. Inhibition of lipoperoxidase activity of the cytochrome c with cardiolipin complex by antioxidants was demonstrated. These findings prove that studies on the effect of antioxidants on this supramolecular nanoparticle will eventually lead to discovery of the news means to prevent and treat diseases caused by apoptosis: myocardial infarction, Alzheimer’s disease, Parkinson’s disease, etc.

## Introduction

Research in the area of programmed cells death, or apoptosis, is moving forward at a rapid rate; this phenomenon is being considered from different perspectives by many natural and biological sciences. Apoptosis was first described in 1972 by J. Kerr, A. Wyllie and A. Currie [1].

A great number of studies is aimed at finding means and methods to manage this phenomenon: either its trigger points or its and inhibition. The research for methods of apoptosis inhibition is crucial for treatment of diseases caused by abnormal cellular “suicide” due to various reasons, such as myocardial infarctions [2], Alzheimer’s disease [3], or, for example, Parkinson’s disease, which most researchers associate with inappropriate apoptosis of brain cells that produce dopamine as a result of oxidative stress [4, 5, 6, 7].

Large number of researches revealed that the cytochrome c–cardiolipin complex is pivotal in the initial stages of apoptosis along the mitochondrial pathway due to its lipoperoxidase activity, which cytochrome c acquires affected by the alteration in its conformation under the exposure to cardiolipin molecules, as a result of which one valence of the iron ion being a part of heme group of protein is released [8, 9, 10]. The primary function of cytochrome c is the transfer of electrons in the mitochondrial respiratory chain [11], in addition, free cytochrome c, once in the cytosol, eventually triggers the caspase cascade of apoptosis reactions [12]. Cytochrome c performs these two functions without neither binding to phospholipids nor acquiring peroxidase activity. Cytochrome c molecule acquires lipoperoxidase activity only after a conformation alteration as a result of the formation of a complex with phospholipids, primarily cardiolipin [8, 9, 10]. Therefore, cytochrome c cannot be considered a classic peroxidase as of horseradish peroxidase, myeloperoxidase, etc. “Facultative peroxidase” is far more appropriate term for cytochrome c, indicating that this protein does not possess peroxidase activity in the native state, but acquires it only under certain conditions. In a living cell, this condition is its binding to cardiolipin.

Kinetics of this lipoperoxidase reaction is assayed via chemiluminescence consisting of recording by a device of superweak luminescence in the visible spectrum, which is prepared from the reactions in the test forming free radicals [13, 14, 15]. However, it should be noted that if a chemiluminescent signal is detected during the study of the properties of the cytochrome *c* with cardiolipin complex, then formally lipoperoxidase activity of this nanoparticle might not be the reason in this case given the process of radical lipid oxidation catalyzed by free iron, known as the Fenton reaction, occurring in the test [16]. Therefore, the objective is being made: to assess the potential role of free (not included into cytochrome *c* heme) iron in chemiluminescence of the cytochrome *c* with cardiolipin complex system. Some researches are being in favor of possible involvement of Fe^2+^ ions in the observed chemiluminescence of the studied system, revealing that cytochrome *c* is destroyed under the exposure of hydrogen peroxide [9] or lipid hydroperoxides [17] and that the effect of some hemoproteins is associated with the release of Fe^2+^ ions from the heme [18]. The primary objective of this study was to assess the potential role of non-heme iron in the chemiluminescence of thecytochrome *c*– cardiolipin system.

Lipoperoxidase activity of intact mitochondria is also advisable to be evaluated in the furtherance of this objective. This is required to prove that the cytochrome c–cardiolipin complex exhibits lipoperoxidase properties as *in vitro*, being a part of the molecular model, but as well inside the living mitochondria.

Due to the fact that apoptosis is the cause of many diseases, it seems crucial to find the method to block it. A very promising direction in this area is the application of antioxidants possessing the mechanism of action employed to inhibit the lipoperoxidase activity of cytochrome c–cardiolipin complex. Dihydroquercetin (as a specie of natural plant antioxidants) and trolox (as a specie of artificial chemically synthesized antioxidants) were selected as antioxidants for the study.

Researches on the effect of antioxidants on the cytochrome c-cardiolipin complex have already been conducted [19, 20], however those works applied luminol as an activator of chemiluminescence, which itself actively participates in chemical reactions with the components of the reaction mixture. It is suggested to apply an activator that enters into such interactions either extremely limited or does not enter at all. As reported by [21, 22, 23], isoquinolysin derivatives of coumarin are being the similar activators, besides being as well lipoperoxidase reactions-specific. Coumarin-334 was applied in this study as a chemiluminescence activator. Coumarin-525 was also applied in the experiment with mitochondria for parallel measurement, as well as also its own chemiluminescence was recorded.

## Materials and methods

The following reagents were applied during experiments: KH_2_PO_4_, 20 mM of buffer solution (pH 7.4); horseradish peroxidase (activity of 112 U/mg, molar mass of 44 173.9 g/mol), 1 mM of aqueous solution prepared from the required sample weight; luminol (5-amino-1,2,3,4-tetrahydro-1,4-phthalazinedione, 3-aminophthalic acid hydrazide), 1 mM of aqueous solution prepared from the required sample weight; peroxide hydrogen, 10 M of aqueous solution (Sigma-Aldrich, USA); antioxidant solutions (trolox, dihydroquercetin), prepared from the required sample weight; coumarin-334, 1 mM of methanol solution, coumarin-525, 1 mM of methanol solution, solutions are prepared from the required sample weight (Sigma-Aldrich, USA); cytochrome C, 1 mM of solution, prepared from the required sample weight; cardiolipin of the bovine heart, 6 mM of methanol solution, prepared from the required sample weight (Avanti Polar Lipids, USA); EDTA solutions, prepared from the required sample weight (Sigma-Aldrich, USA); ortho-phenanthroline (o-phenanthroline) solutions, prepared from the required sample weight (REAKHIM, Russia); 5 mM of copperas solution, prepared from the FeSO_4_∙7H_2_O sample of required weight (REAKHIM, Russia); soy lecithin, a solution with concentration of 10 mg/mL, prepared from the required sample weight.

Solutions with the required for experiments concentrations were obtained via sequential dilution method, one reciprocal dilution was maximum 10.

### Chemiluminescent methods of study

Studies on the chemiluminescence measurement were performed on “Lum-5773” chemiluminometer made by LLC “DISoft” (Russia), connected to a PC with “PowerGraph” software. Chemiluminometer was calibrated on uranium glass prior to each series of measurements.

#### Effect of o-phenanthroline and EDTA on chemiluminescence in the Fe^2+^ions–soy lecithin system

50 µL of 2 mM of copperas (FeSO_4_) and 50 µL of ortho-phenanthroline or EDTA solution of various concentrations were added into the empty flow cell followed by the cell insertion into the chemiluminometer cell holder and initiation of chemiluminescence recording. In 30 seconds, 800 µL of a solution containing 25 µL of 1 mM coumarin-334 methanol solution and 775 µL of 20 mM phosphate buffer was added into the chemiluminometer flow cell. By the 60^th^ second of chemiluminescence recording, 200 µL of soy lecithin solution in a concentration of 10 mg/mL was added followed by luminescence recording for the required amount of time.

#### Effect of o-phenanthroline and EDTA on chemiluminescence in the horseradish peroxidase–soy lecithin system

50 µL of 210 µM of horseradish peroxidase and 50 µl of ortho-phenanthroline or EDTA solutions in various concentrations were added into the flow cell followed by the cell insertion into the chemiluminometer cell holder and initiation of chemiluminescence recording. In 30 seconds, 800 µL of a solution containing 25 µL of 1 mM coumarin-334 methanol solution and 775 µL of 20 mM phosphate buffer was added into the chemiluminometer flow cell. By the 60^th^ second of chemiluminescence recording, 200 µL of soy lecithin solution in a concentration of 10 mg/mL was added followed by luminescence recording for the required amount of time.

#### Effect of o-phenanthroline and EDTA on chemiluminescence in the horseradish peroxidase – bovine cardiolipin system

50 µL of 210 µM of cytochrome c and 50 µL of ortho-phenanthroline or EDTA solutions in various concentrations were added into the flow cell followed by the cell insertion into the chemiluminometer cell holder and initiation of chemiluminescence recording. In 30 seconds, 800 µL of a solution containing 25 µL of 1 mM coumarin-334 methanol solution and 775 µL of 20 mM phosphate buffer was added into the chemiluminometer flow cell. By the 60^th^ second of chemiluminescence recording, 100 µL of 6.6 mM bovine cardiolipin was added followed by luminescence recording for the required amount of time.

#### Effect of antioxidants on lipoperoxidative activity of the cytochrome c– cardiolipin complex

100 µL of 100 µM cytochrome *c* and 50 µL of trolox or dihydroquercetin solutions in various concentrations were added into the empty flow cell followed by the cell insertion into the chemiluminometer cell holder and initiation of chemiluminescence signal recording. In 30 seconds, 800 µL of a solution containing 25 µL of 1 mM coumarin-334 methanol solution and 775 µL of 20 mM phosphate buffer was added into the chemiluminometer flow cell. At the 60^th^ second of chemiluminescence recording, 100 µL of 6 mM of bovine cardiolipin was added followed by chemiluminescence recording for 250 seconds. Subseqently, 100 µL of 1 mM H_2_O_2_ was added into the flow cell and inserted into the cell holder and, at the 360th second after the measurement initiation, 800 µL of contents of the first cell was transferred into this cell followed by luminescence recording for the required amount of time.

### Study of mitochondrial suspension chemiluminescence induced by hydrogen peroxide with sodium nitride (azide)

920 µL of mitochondrial suspension isolated by below-described method was added into the empty chemiluminometer flow cell. In about 230 seconds, 800 µL of a solution containing 25 µL of 1 mM coumarin-334 or coumarin-525 methanol solution or 25 µL of methanol was added into the flow cell (upon measurement of own chemiluminescence); by approx. 500^th^ second of chemiluminescence recording, 5 µL of hydrogen peroxide was added; by the 750^th^ second, 50 µL of 2 mM sodium nitride was added followed by addition of another 5 µL of hydrogen peroxide by 900^th^ second of chemiluminescence recording with subsequent luminescence recording for the required amount of time.

#### Isolation of rat liver mitochondria

Mitochondria were isolated from the liver of male *Wistar* rats (275±25 g). Euthanasia was performed by asphyxiation of a rat with carbon dioxide. A liver sample weighing about 4 g was washed in an isolation buffer (10 mM HEPES, 1 mM EDTA, 70 mM sucrose and 200 mM mannitol, pH 7.5). The sample was homogenized in a 40 mL separation buffer containing 1 mg/mL of bovine serum albumin which was purified from lipids. The final suspension was centrifuged at 600 G for 5 minutes. Supernatant was separated and centrifuged at 11 000 G for 10 minutes, after which the supernatant was removed and the precipitate was resuspended in 40 mL of separation buffer. The final suspension was centrifuged at 600 G for 5 minutes. Supernatant was separated and centrifuged at 11 000 G for 10 minutes followed by the supernatant removal, the precipitate was resuspended in a storage buffer (10 mM HEPES, 250 mM sucrose, 1 mM ATP, 0.08 mM ADP, 5 mM sodium succinate and 2 mM KH_2_PO_4_, pH 7.5) at 0-4°C on ice.

### Findings

#### Evaluation of the potential role for non-heme iron in chemiluminescence of the cytochrome c-cardiolipin complex

Potential role for non-heme iron in the chemiluminescence of the cytochrome c–cardiolipin system was studied to ensure the reliability of findings on the peroxidase activity of the cytochrome c with cardiolipin complex obtained by measuring chemiluminescence. For this purpose, the chemiluminescence of three systems (soy lecithin-Fe^2+^, soy lecithin – horseradish peroxidase, cytochrome c– cardiolipin) and the effect of complexon addition on iron (o-phenanthroline and ethylenediaminetetraacetate (EDTA)) of various concentrations was recorded. Lipid substrate was added into a flow cell with coumarin-334, complexon and Fe^2+^ ions or protein by 60^th^ second after the measurement initiation throughout all of the experiments, described in this Chapter.

Diagrams in figure 1 clearly demonstrate that o-phenanthroline (Fig. 1A) and EDTA (Fig. 1B) inhibit the Fe^2+^-induced chemiluminescence. Subsequently, a similar effect of complexones was studied, but this time on the soy lecithin– horseradish peroxidase system.

**Fig. 1.**
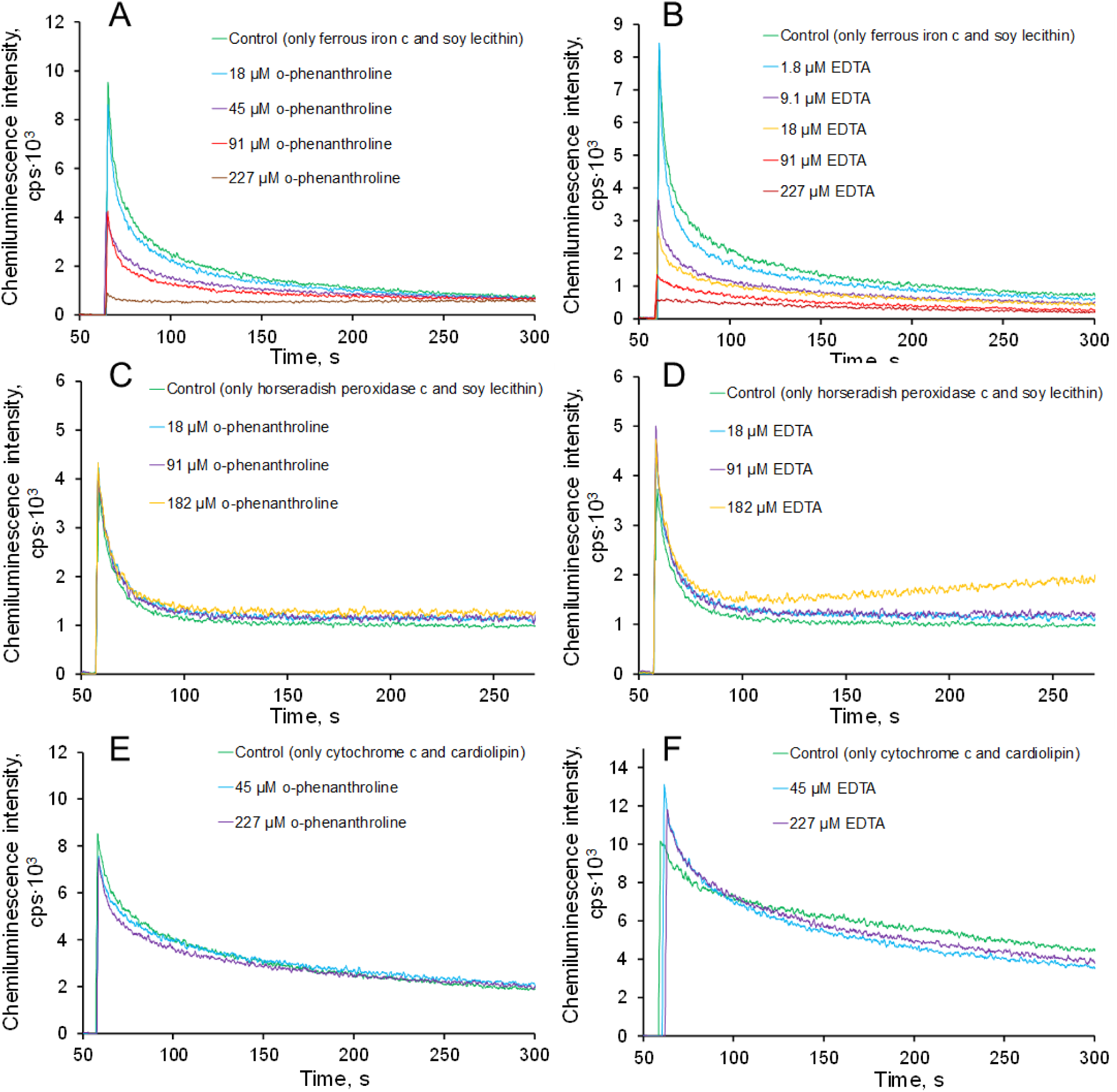
**A-B.** Effect of o-phenanthroline (A) and EDTA (B) on Fe^2+^-induced chemiluminescence kinetics with soy lecithin. FeSO_4_– 91 µm, lecithin – 2 mg/mL, o-phenanthroline or EDTA (concentrations are given in the figure) **C-D.** Effect of o-phenanthroline (C) and EDTA (D) on the chemiluminescence kinetics with horseradish peroxidase and soy lecithin. Horseradish peroxidase – 10 µM, soy lecithin – 2 mg/mL, o-phenanthroline or EDTA (concentrations are given in the figure). **E-F.** Effect of o-phenanthroline (E) and EDTA (F) on the chemiluminescence kinetics with cytochrome c and bovine cardiolipin. Cytochrome c – 10 µM, bovine cardiolipin – 600 µM, o-phenanthroline or EDTA (concentrations are given in the figure).

Findings represented in figure 1C-D indicates that neither o-phenanthroline nor EDTA inhibit the chemiluminescence in the soy lecithin–horseradish peroxidase system, shape of the curve is generally similar to that of the soy lecithin–Fe^2+^ system.

Next, the effect of o-phenanthroline and EDTA on the chemiluminescence in the cytochrome c–cardiolipin system was studied. Diagrams in figure 1E-F prove that no significant decrease of the chemiluminescence intensity by o-phenanthroline and EDTA effect in the cytochrome c–cardiolipin system was detected.

#### Study of lipoperoxidase activity in isolated mitochondria of Rattus Norvegicus Wistar liver

Researches of lipoperoxidase activity of the cytochrome c–cardiolipin complex were conducted on molecular models, *in vitro*. However, that system is generally artificial; therefore, it is required to prove the possible processes revealed on molecular models are occurring in intact mitochondria, meaning that the suspension of intact mitochondria with hydrogen peroxide is capable of chemiluminescence. For this purpose, chemiluminescence of an isolated mitochondrial suspension from *Rattus norvegicus Wistar* rat liver cells with addition of hydrogen peroxide was measured, indicating an available component in mitochondria possessing the peroxidase activity. This component is thought to be the cytochrome c-cardiolipin complex.

The experiment findings are given in figure 2. After the first addition of hydrogen peroxide, a fairly rapidly subsiding flash is noted, whereas its amplitude in coumarin-334 or coumarin-525 test (black curves) is higher than in the test without these derivative, meaning that this flash is caused by lipid radicals formation (gray curve), since coumarin-334 and coumarin-525 are specific activators on them. After the rapid subside of this flash, sodium azide was added followed by addition of the same amount of hydrogen peroxide as was added the first time. This time the amplitude of the chemiluminescence flash was higher in comparison to azide-absence in the system. This flash lasted far longer. Test without coumarin (gray curve), also revealed a long-lasting flash, however of lower intensity due to the fact that it was the system’s own chemiluminescence.

**Fig. 2.**
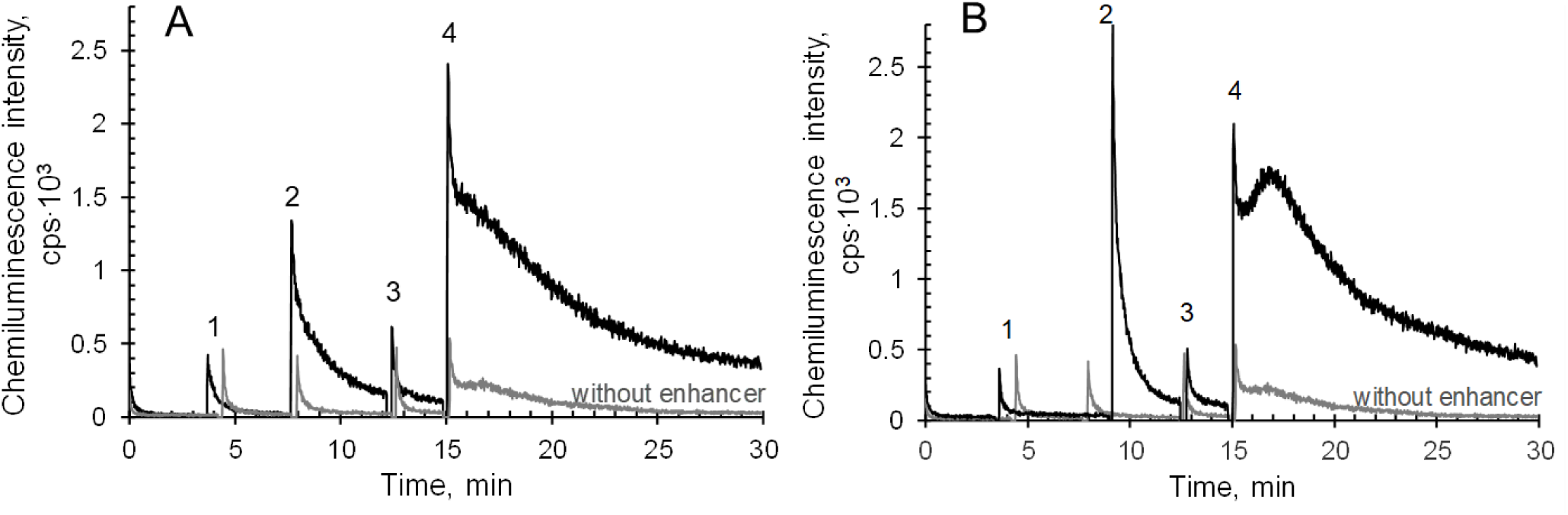
H_2_O_2_-induced mitochondrial chemiluminescence exposed to sodium azide, activated by coumarin-334 (A) or coumarin-525 (B), gray curve – methanol was added instead of the activator. Numbers indicate additions to the cell: 1 – 920 µL of mitochondria, 2 – 25 µL of 1 mM coumarin-334 (A) or coumarin-525 (B) or 25 µL of methanol for control (gray curve), 3 – 5 µL of 96 mM H_2_O_2_, 4 – 50 µL of 2 mM NaN_3_, 5 – 5 µL of 96 mM H_2_O_2_; concentrations of substances in the system: coumarin-334 (A) or coumarin-525 (B) – 25 µM, sodium azide – 100 µM, H_2_O_2_ (after the first addition) – 480 µM.

Chemiluminescence in the case of coumarin-525 (Fig. 2B) was very similar to that of coumarin-334 (Fig. 2A): lengthening of the flash was also observed with hydrogen peroxide, affected by presence of sodium azide in the system, inhibiting the actions of mitochondrial catalases. This led to increase of peroxide concentration in the system (in the case of the first flash, most of the hydrogen peroxide was processed by catalases). And this resulted in generation of the large amount of lipid radicals due to the lipoperoxidase activity of mitochondria affected by present cytochrome c–cardiolipin complex.

The difference in shape of chemiluminescence curves with coumarin-334 and coumarin-525 is probably due to the fact that coumarin-525, unlike coumarin-334, may exhibit an antioxidant effect given the benzimidazole group available in its structure [24].

#### Effect of antioxidants on lipoperoxidative activity of the cytochrome c– cardiolipin complex

In the course of this work, the effect of dihydroquercetin (taxifolin) and trolox on the kinetics of the free radical formation reaction in the cytochrome c– cardiolipin complex – H_2_O_2_ system was also studied. Assessment of this formation was conducted via chemiluminescence: the dependence of the signal intensity on the reaction time of the interaction of the components of the reaction mixture was detected – the higher the signal intensity, the higher the rate of free radicals formation in the system at this point of time.

Effect of dihydroquercetin on the chemiluminescence of the given system should be reviewed in more details.

Diagrams in figure 3 clearly indicate the effect of dihydroquercetin on the reaction with hydrogen peroxide: an acute chemiluminescence inhibition, and hence the free radicals formation in comparison with the control test, and this inhibition is stronger as higher the concentration of dihydroquercetin.

**Fig. 3.**
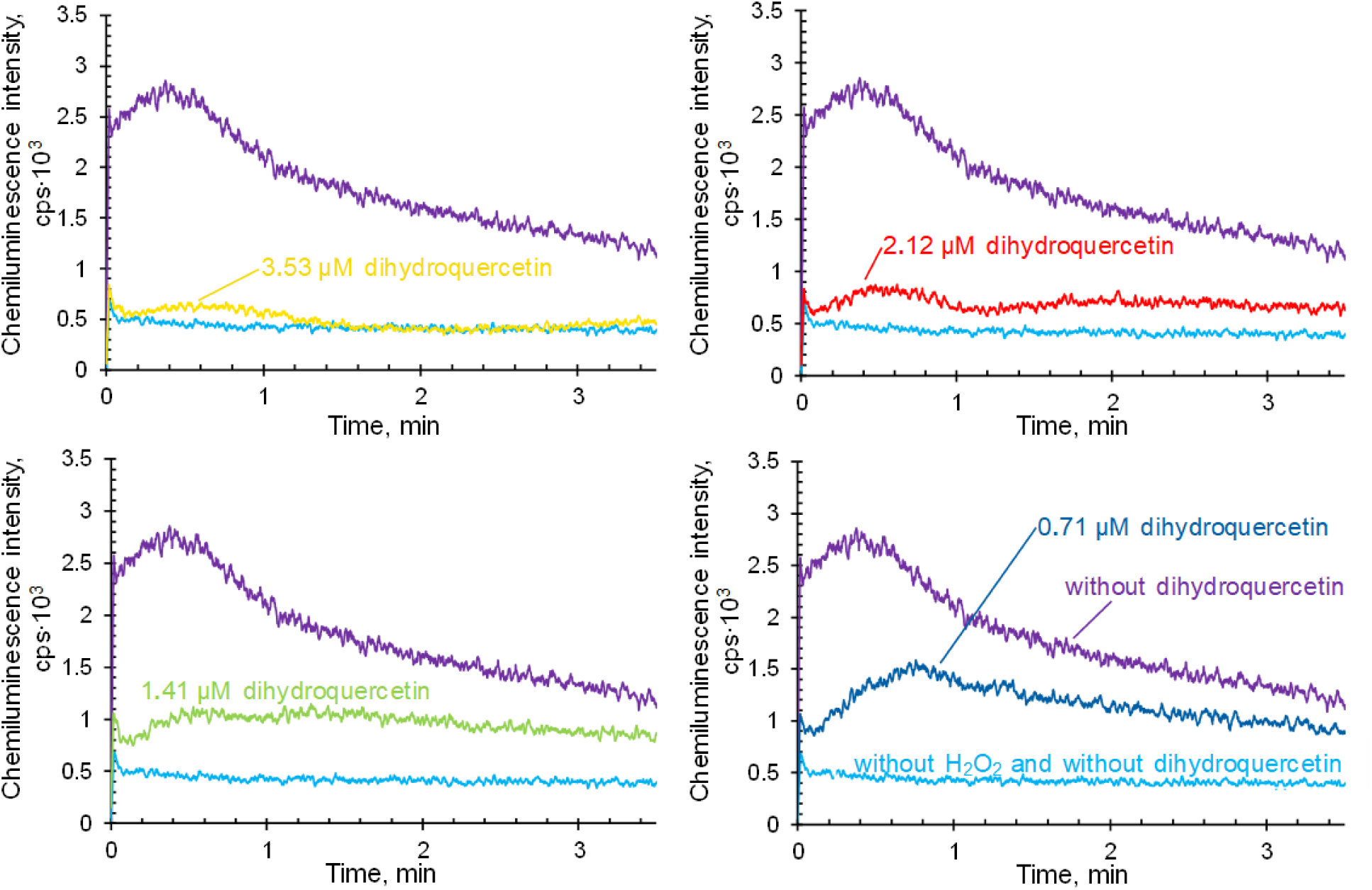
Ihibition of lipoperoxidase activity of the cytochrome c-cardiolipin complex by dihydroquercetin. Cytochrome c – 10 µM, bovine cardiolipin – 600 µM, H_2_O_2_ – 500 µM, coumarin-334 – 25 µM. Light blue curves– control without hydrogen peroxide, purple curves– without dihydroquercetin. The remaining curves – chemiluminescence at various concentrations of dihydroquercetin.

5 measurements were conducted for each concentration of dihydroquercetin. The average maximum chemiluminescence intensity (amplitude) for each dihydroquercetin concentration was estimated.

The dose-dependent inhibition of dihydroquercetin by hydrogen peroxide-induced chemiluminescence is clearly identified. 50% inhibition of peroxidase activity is observed at a concentration of dihydroquercetin equal to ≈0.55 µM (Fig. 4), with a ratio of bovine cardiolipin: cytochrome c 60:1. The authors [20] in a similar experiment with the ratio of bovine cardiolipin: cytochrome c 32:1 have obtained a half quenching concentration of dihydroquercetin equal to 10 µm.

**Fig. 4.**
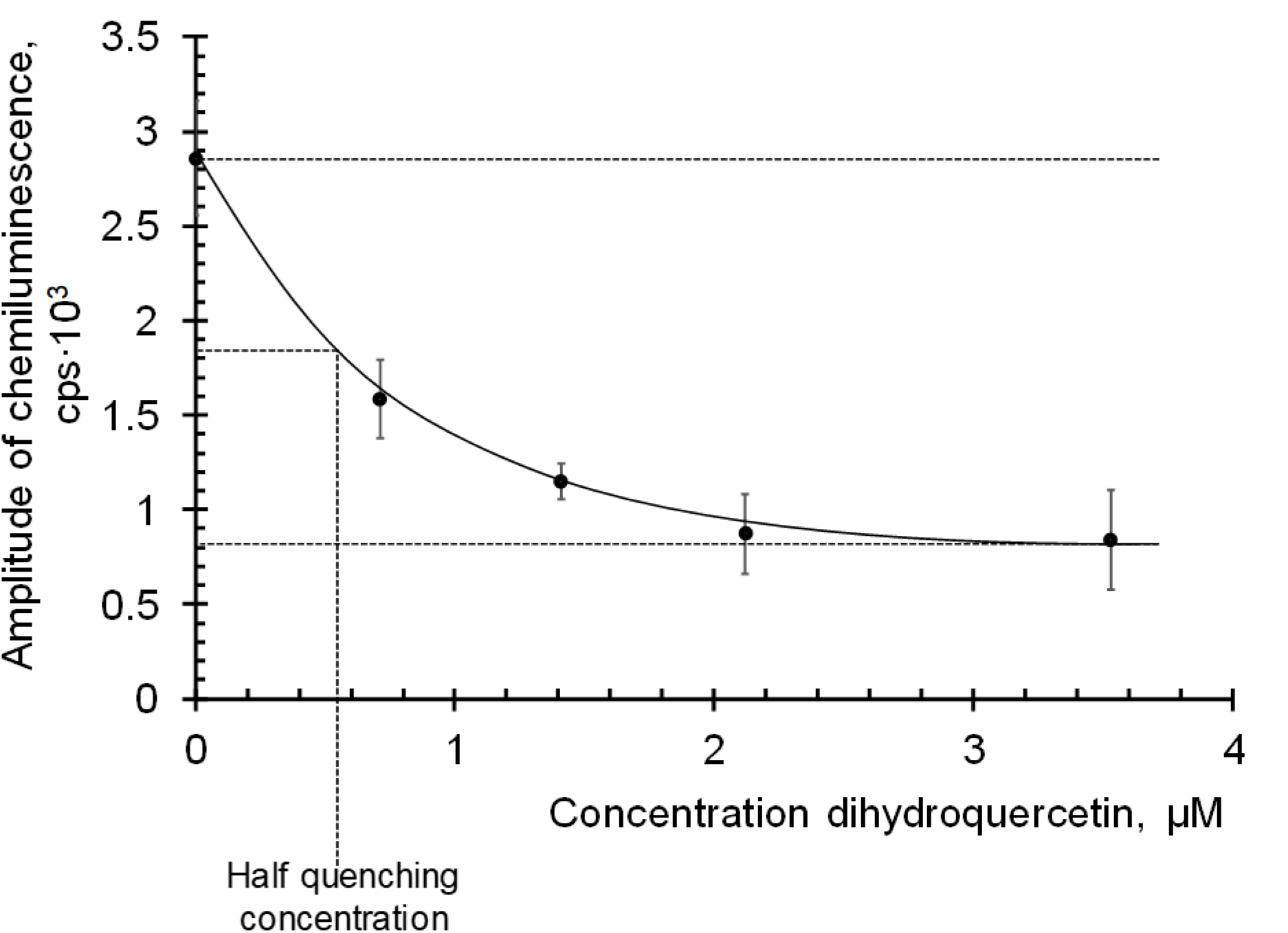
Relation of chemiluminescence amplitude to concentration of dihydroquercetin.

The effect of another antioxidant – trolox on the hydrogen peroxide-induced chemiluminescence was also studied.

Diagrams in figure 5 demonstate that trolox effect somewhat differs from the dihydroquercetin effect: in general, no absolute inhibition of the chemiluminescence intensity is observed, however a delay in time of a slow flash is clearly exhibited, and this delay is greater as higher the trolox concentration.

**Fig. 5.**
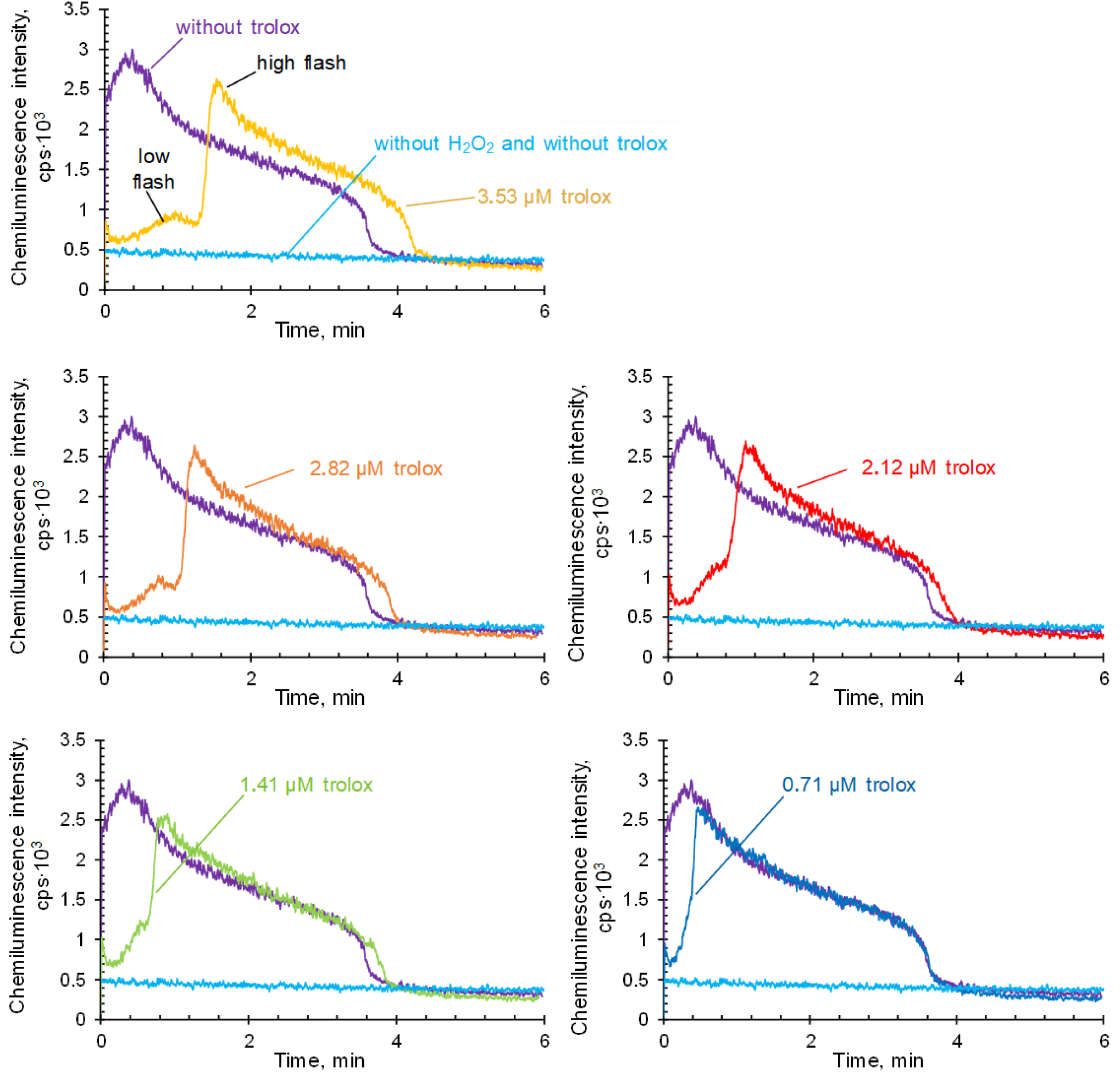
Effect of trolox on lipoperoxidase function of the cytochrome c– cardiolipin complex. Cytochrome c – 10 µM, bovine cardiolipin – 600 µM, H_2_O_2_ – 500 µM, coumarin-334 – 25 µM. Light blue curves – control without hydrogen peroxide, purple curves – without trolox. The remaining curves are chemiluminescence at various concentrations of trolox in the system.

No significant dependence of the high flash amplitude to the concentration of trolox was found. Notably the chemiluminescence flash develops in two stages: at first a low intensity flash was observed followed by a high intensity flash. Both flashes, low and high, are dose-dependently delayed by trolox, and the intensity of both flashes does not depend on the concentration of trolox, however is somewhat lower in comparison with the test without an antioxidant (blue curve in diagrams of figure 5).

## Discussion

### Evaluation of the potential role for non-heme iron in chemiluminescence of the cytochrome c-cardiolipin complex

The suspicion that the observed chemiluminescence in the cytochrome c– cardiolipin system may be due to Fe^2+^ ions occurred in this case due to the fact that the chemiluminescence kinetics in the presence of cytochrome c–cardiolipin complex was similar to the kinetics obtained earlier in the system with Fe^2+^ ions-triggered chemiluminescence [25]. Besides, some researches are being in favor of possible involvement of Fe^2+^ ions in the observed chemiluminescence of the studied system, revealing that cytochrome c is destroyed under the exposure of hydrogen peroxide [9] and lipid hydroperoxides [17] and that the effect of some hemoproteins is associated with the release of Fe^2+^ ions from the heme [18].

Potential role of non–heme iron in chemiluminescence observed in the cytochrome c–cardiolipin system was studied herein by comparing the means of iron complexons (o–phenanthroline and EDTA) affecting chemiluminescence with lipid substrate (soy lecithin) triggered by Fe^2+^ or horseradish peroxidase ions over the effect of the same complexons on chemiluminescence observed in the cytochrome c–cardiolipin system. It was assumed that the complexons would inhibit Fe^2+^ ions-triggered chemiluminescence without inhibiting horseradish peroxidase-triggered chemiluminescence.

It should be noted that concentrations of horseradish peroxidase and cytochrome c were the same (10 µM), but the concentration of Fe^2+^ was 91 µM. The selection of such concentration of iron was considering the fact that iron ions of such concentration generated the same chemiluminescence amplitude as the mentioned proteins at a concentration of 10 µM. This implies, that the lipoxygenase effect of free Fe^2+^ ions is almost 10 times less than that of peroxidases, and since horseradish peroxidase is a classical peroxidase, thus the similarity of chemiluminescence of cytochrome c–cardiolipin and horseradish peroxidase soy–lecithin systems can indeed be considered the proof of cytochrome c–cardiolipin complex being functionally a peroxidase.

Reviewing the chemiluminescence curves in figure 1, it may be noted that both EDTA and o-phenanthroline inhibit Fe^2+^-induced chemiluminescence, and this inhibition is greater as higher the complexon concentration. At the same time, these complexons, derived at the same concentrations, did not inhibit horseradish peroxidase-induced chemiluminescence. Horseradish peroxidase is being considered a classical peroxidase, therefore the chemiluminescence in its presence is believed to be due to the involvement of peroxidase enzyme instead of Fe^2+^ ions. The striking difference in complexons effect on the two described chemiluminescent systems allows to distinguish Fe^2+^-induced chemiluminescence from chemiluminescence with peroxidases, which in this case offers a solution to what causes chemiluminescence observed in the cytochrome c–cardiolipin system. Obtained findings indicates that the complexons effect the chemiluminescence in the cytochrome c–cardiolipin system in the same way as chemiluminescence with the classical peroxidase, and that this effect is strikingly differs from the means these complexons manifest in the system with Fe^2+^ ions. Therefore, the conclusion is that the chemiluminescence of the cytochrome c–cardiolipin system is implicated by peroxidase enzyme activity rather than of free Fe^2+^ ions appearing in the system after the destruction of the cytochrome c preparation, for example, during storage.

### Study of lipoperoxidase activity in isolated mitochondria of Rattus Norvegicus Wistar liver

The goad objective was to assay whether mitochondria have lipoperoxidase activity, described herein on the molecular model – cytochrome c–cardiolipin system, meaning that the *in vitro* effects described herein of cytochrome c– cardiolipin complex as well occur *in vivo*.

Lipoxygenase activity is manifested in chemiluminescence without hydrogen peroxide in the system or at its low concentrations, lipoperoxidase activity in the other hand -in chemiluminescence at high concentrations of H_2_O_2_ [26]. It is worth noting that the presence of inactivated mitochondrial chemiluminescence was discovered in the 1960s by Y.A. Vladimirov and O.F. Lvova, where the intensity of the luminescence correlated with the mitochondrial capacity to oxidative phosphorylation [13, 27, 28, 29, 30, 31]. A possible reason for the inactivated mitochondrial chemiluminescence was the lack of enzymes for β-oxidation of fatty acids, leading to the direct oxidation of fatty acids by molecular oxygen followed by the formation of lipid peroxides with their subsequent emission of quanta of light [28]. But it is also possible that this chemiluminescence is the effect of lipoxygenase activity of the cytochrome c– cardiolipin complex retained in mitochondria.

Coumarin-334 or coumarin-525-activated chemiluminescence or own mitochondrial suspension chemiluminescence was recorded during this study. Luminescence prior to the addition of hydrogen peroxide indicates the lipoxygenase activity of mitochondria, a mild chemiluminescence intensity was observed within the background values. Instead, clearly visible flashes from the addition of hydrogen peroxide was noted given the lipoperoxidase activity of mitochondria.

The conclusion is that sodium azide facilitates the leveling of factors removing hydrogen peroxide from the system since the flash after the first addition of hydrogen peroxide quickly subsided, and after the second one, conducted after the addition of sodium azide to the system, had a longer duration resulting in appearance of a higher steady-state condition compared to the initial background (Fig. 2). Most likely, sodium azide, as indicated in [32], inhibited the activity of catalases retained in mitochondria, so that other components of mitochondria, which is thought to be the cytochrome c–cardiolipin complex, were able to fully exhibit lipoperoxidase activity, leading to luminescence of quanta of light via pathways described by many different authors and collected in one edition [14].

Thus, lipoperoxidase activity was first established with application of mitochondria from rat liver and chemiluminescence was observed without hydrogen peroxide, allowing the conclusion that mitochondria possess lipoperoxidase activity, while sodium nitride (azide) significantly enhanced lipoperoxidase function, which is most likely due to the inhibition of catalase activity.

### Effect of antioxidants on lipoperoxidative activity of the cytochrome c– cardiolipin complex

Studies on the effect of antioxidants on the cytochrome *c* with cardiolipin complex have already been conducted [19, 20, 33]. However, the increase of quantum efficiency of chemiluminescence in them was done by application of either luminol [20, 33], entering into direct chemical interaction with the components of the studied system, or coumarin-525 [20], which potentially may exhibit an antioxidant effect [24] due to purine arrangement retained in its structure, which may cause erroneous results. Coumarin-334 was selected for this study as an activator of chemiluminescence considering the absence of benzimidazole group in its structure.

As previously mentioned, trolox and dihydroquercetin (taxifolin) were applied as antioxidants, their formulas are given in figure 6.

**Fig. 6.**
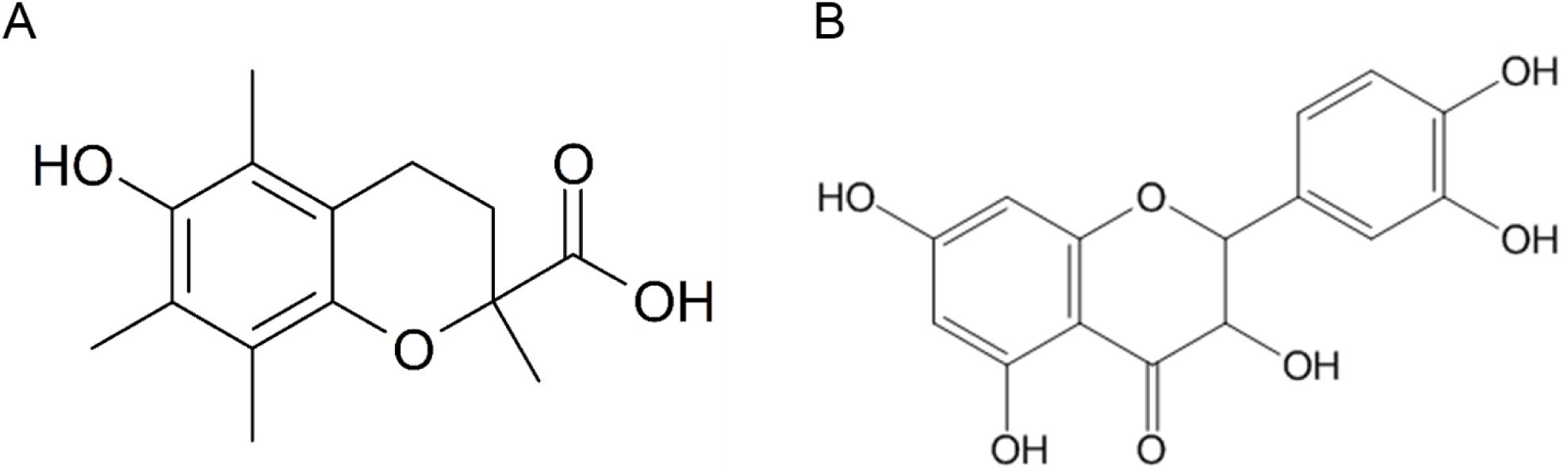
Formulas of trolox (A) and dihydroquercetin (B)

Trolox in concentrations from 2.12 µM to 3.53 µM dose-dependently delays the development of the chemiluminescence flash. A low-amplitude (low) chemiluminescence flash occurs before a sharp rise of the curve and a maximum of chemiluminescence (high flash) at trolox concentrations from 2.12 µM. A high flash is delayed for a shorter period of time, and the low flash can no longer be clearly differentiated exposed to trolox concentrations below 2.12 µM. A brief latency period should be noted occurring before the low flash (chemiluminescence curves in Fig. 5).

The multistage of chemiluminescence kinetics may be a consequence of two hydroxyl groups available in trolox, potentially possessing antioxidant properties, and, most likely, independently of each other, one hydroxyl is exposed at the first stage of the reaction, and the second one at the other stage of the reaction.

Concerning dihydroquercetin containing five phenolic hydroxyl, the following is determined: there is no latency period (chemiluminescence curves -in Fig. 3), no distinct dependence of the time of maximum chemiluminescence flashes onset on dihydroquercetin concentration. However, their amplitude clearly depends on dihydroquercetin concentration (Fig. 3), whereas trolox does not possess the similar dependence, what is more it that the chemiluminescence intensity does not reach the control level within 3.5 minutes.

This findings on dose–dependent chemiluminescence inhibition, and hence the peroxidase activity of cytochrome c–cardiolipin complex along with free radicals formation are compatible with those available in the literature: so, inhibition in a similar system was described in [20] back in 2008, as well as inhibition of chemiluminescence on the antioxidant-depending concentration was described in [26, 33].

Diagram in figure 4 clearly indicates the dose-dependent inhibition of dihydroquercetin by hydrogen peroxide-induced chemiluminescence. 50% inhibition of peroxidase activity is observed at a concentration of dihydroquercetin equal to 0.51 µM, with a ratio of bovine cardiolipin: cytochrome *c* 60:1. The authors [20] in a similar experiment with the ratio of bovine cardiolipin: cytochrome *c* 32:1 have obtained a dihydroquercetin concentration of half attenuation equal to 10 µM.

Also, the article [33] provides description of a study of the effect of antioxidants on the peroxidase activity of cytochrome c– 1,1’,2,2’-tetraoleylcardiolipin complex, for trolox and dihydroquercetin concentrations of half quenching chemiluminescence were established as follows: 3.7 µM and 0.7 µM respectively. It should be also pointed out that chemiluminescence in the above-mentioned study was activated by luminol [33], directly chemically interacting with system components. Noteworthy the fact that the chemiluminescence curves, given by the authors of the article [33], in their forms do not correspond to those obtained herein for either dihydroquercetin or trolox. Most likely it does not depend from the various ratio of bovine cardiolipin: cytochrome c applied for some experiments herein or that tetraoleylcardiolipin was applied in others but given the chemiluminescence activation was conducted not by a physical activator, as was done herein with coumarin-334 application, but by luminol probe-induced chemiluminescence, chemically interacting with components of the system.

Thus, the inhibition of lipoperoxidase activity of cytochrome c–cardiolipin complex by dihydroquercetin and its temporary inhibition by trolox was established enabling more detailed studies of these antioxidants in terms of their application for prevention of apoptosis onset for vital tissue cells.

## Conclusion

Coumarin-334-activated chemiluminescence was applied in order to record lipoperoxidase and lipoxygenase activity of cytochrome c–cardiolipin complex, moreover, the chemiluminescence accompanying lipoxygenase activity is proved to be caused by the activity of this supramolecular nanoparticle rather than Fenton reaction due to the presence of free, non-heme iron in the system: these complexons inhibited chemiluminescence in the soy lecithin-Fe^2+^ system, but no inhibition of chemiluminescence in the cytochrome c–cardiolipin system was detected.

Dose–dependent inhibition of lipoperoxidase activity of the cytochrome c-cardiolipin complex by trolox and dihydroquercetin (taxifolin) was identified, meaning the possible application of antioxidants in order to inhibit this process in living cells, therefore to block apoptosis and prevent, and perhaps cure, many neurodegenerative diseases, strokes, etc.

Applicability of herein obtained findings to living systems via detecting the lipoperoxidase activity in intact, living mitochondria was proven suggesting that the properties of assayed herein the cytochrome c–cardiolipin complex and described in previous studies, are inherent *in vivo* as well as *in vitro*. Prospective research may continue the study on living systems: in particular, study the effect of antioxidants on lipoperoxidative activity of mitochondria with inhibited catalases, subsequently the research on cultures of cells should be continued followed by the study on tissues, and potentially transit to the study of aspects established herein on laboratory animals and initiate the research for methods to inhibit lipoperoxidative and lipoxygenase activity of the cytochrome c–cardiolipin complex *in vivo* aiming at apoptosis inhibition, prevention and treatment of diseases caused by death of cells, pivotal for the normal functioning of the human body.

## Summary

1. The chemiluminescence of the cytochrome c–cardiolipin complex – hydrogen peroxide system was determined to be generated by lipoxygenase and lipoperoxidase activity of the cytochrome c-cardiolipin complex rather than non-heme iron.
2. Lipoperoxidase activity of mitochondria was first established via application of mitochondria from rat liver (*Rattus Norvegicus Wistar rat*), whereas, sodium nitride (azide) significantly increased the lipid radicals formation indicating the connection with inhibition of mitochondrial catalases activity.
3. Effect of trolox and dihydroquercetin (taxifolin) antioxidants on lipoperoxidase activity of the cytochrome c–cardiolipin complex was studied. Dose–dependent inhibition of lipoperoxidase activity of the cytochrome c-cardiolipin complex by dihydroquercetin and its temporary inhibition by trolox (onset of the latency period) was established.

## Author contributions

L.A. Romodin has designed the experimentations, conducted the test, processed the findings, wrote the text of the article, drawn up the illustrations, Y.A. Vladimirov has suggested the concept, designed the experimentations, N.P. Lysenko has done administrative works, V.B. Chernetsov has financed the publication of the article, its translation into English, Y.V. Antonova has supervised the translation or has done the translation of the article into English, has managed the correspondence with the editors of the magazine.

## Acknowledgements

We thank our colleague Mr G.K.Vladimirov for managing the research and Ms Zarudnaya E.N. for consultations on biochemistry.

